# A large-scale retrospective study in metastatic breast cancer patients using circulating tumor DNA and machine learning to predict treatment outcome and progression-free survival

**DOI:** 10.1101/2023.03.03.530936

**Authors:** Emma J Beddowes, Mario Ortega-Duran, Solon Karapanagiotis, Alistair Martin, Meiling Gao, Riccardo Masina, Ramona Woitek, James Tanner, Fleur Tippin, Justine Kane, Jonathan Lay, Anja Brouwer, Stephen-John Sammut, Suet-Feung Chin, Davina Gale, Dana Tsui, Sarah Jane Dawson, Nitzan Rosenfeld, Maurizio Callari, Oscar M Rueda, Carlos Caldas

## Abstract

**Purpose:** Monitoring levels of circulating tumor-derived DNA (ctDNA) represents a non-invasive snapshot of tumor burden and potentially clonal evolution. Here we describe how a novel statistical model that uses serial ctDNA measurements from shallow whole genome sequencing (sWGS) in metastatic breast cancer patients produces a rapid and inexpensive assessment that is predictive of treatment response and progression-free survival.

**Patients and Methods:** A cohort of 188 metastatic breast cancer patients had DNA extracted from serial plasma samples (total 1098, median=4, mean=5.87). Plasma DNA was assessed using sWGS and the tumor fraction in total cell free DNA estimated using ichorCNA. This approach was compared with ctDNA targeted sequencing and serial CA 15-3 measurements. The longitudinal ichorCNA values were used to develop a Bayesian learning model to predict subsequent treatment response.

**Results:** We identified a transition point of 7% estimated tumor fraction to stratify patients into different categories of progression risk using ichorCNA estimates and a time-dependent Cox model, validated across different breast cancer subtypes and treatments, outperforming the alternative methods. We then developed a Bayesian learning model to predict subsequent treatment response with a sensitivity of 0.75 and a specificity of 0.66.

**Conclusion:** In patients with metastatic breast cancer, sWGS of ctDNA and ichorCNA provide prognostic and predictive real-time valuable information on treatment response across subtypes and therapies. A prospective large-scale clinical trial to evaluate clinical benefit of early treatment changes based on ctDNA levels is now warranted.

## INTRODUCTION

Breast cancer is the most common cancer diagnosis and the fifth leading cause of cancer death worldwide. Treatment options for patients with metastatic breast cancer have greatly increased but there remains an unmet need to monitor therapy response in real time^1–3^. Accurate real-time methods of monitoring treatment response are required to minimize time spent on ineffective therapies and improve access to more effective therapy. CA15-3, a tumor marker available in the clinic is often used to monitor response but has limited sensitivity and dynamic range^4^. We have shown it has inferior performance when compared with circulating tumor DNA (ctDNA)^4^. ctDNA assays can also provide a rapid, non-invasive and dynamic way of tracking genomic evolution and detecting the emergence of resistance mutations which could prompt therapy change^5–7^.

Breast cancer genomic landscapes are dominated by chromosomal copy number aberrations (CNAs), with around 85% of tumor gene expression changes driven by these CNAs ^8–10^. CNAs can be profiled using shallow whole genome sequencing (sWGS) of plasma DNA as a rapid and cheap method to characterize CNAs in ctDNA. Crucially, the detection of ctDNA in plasma using sWGS does not rely on any prior knowledge of the originating tumor genome.

Here, we evaluated the utility of ctDNA quantification using sWGS to predict treatment response in a consecutive cohort of metastatic breast cancer patients. We assessed the performance of established analysis tools to measure ctDNA levels, including ichorCNA^11^, z-score^12^ and t-MAD^13^ and developed a Bayesian learning model that uses data from serial ctDNA measurements to dynamically predict treatment response. We also compared this approach to the use of a ctDNA targeted sequencing panel and measuring CA15-3 in the same plasma samples to determine how these different methods performed in predicting treatment response.

## PATIENTS AND METHODS

### Patient Cohort and Sample Collection

A cohort of 188 patients with metastatic breast cancer was recruited into the DETECT clinical study at Cambridge University Hospitals (CUH), UK between 2012-2019. Eligible patients were those women with metastatic breast cancer undergoing treatment. Serial blood samples were collected at specific time points as shown in Figure 1. For patients on chemotherapy, blood samples were taken prior to the next cycle of therapy and for a minimum of four cycles. For patients on continuous treatments such as endocrine therapy, blood samples were taken at routine clinic visits (typically every 3-6 months). Cohort composition for each analysis is shown in Figure 1a-c and the REMARK table is shown in the Supplementary Methods.

**FIGURE 1.**
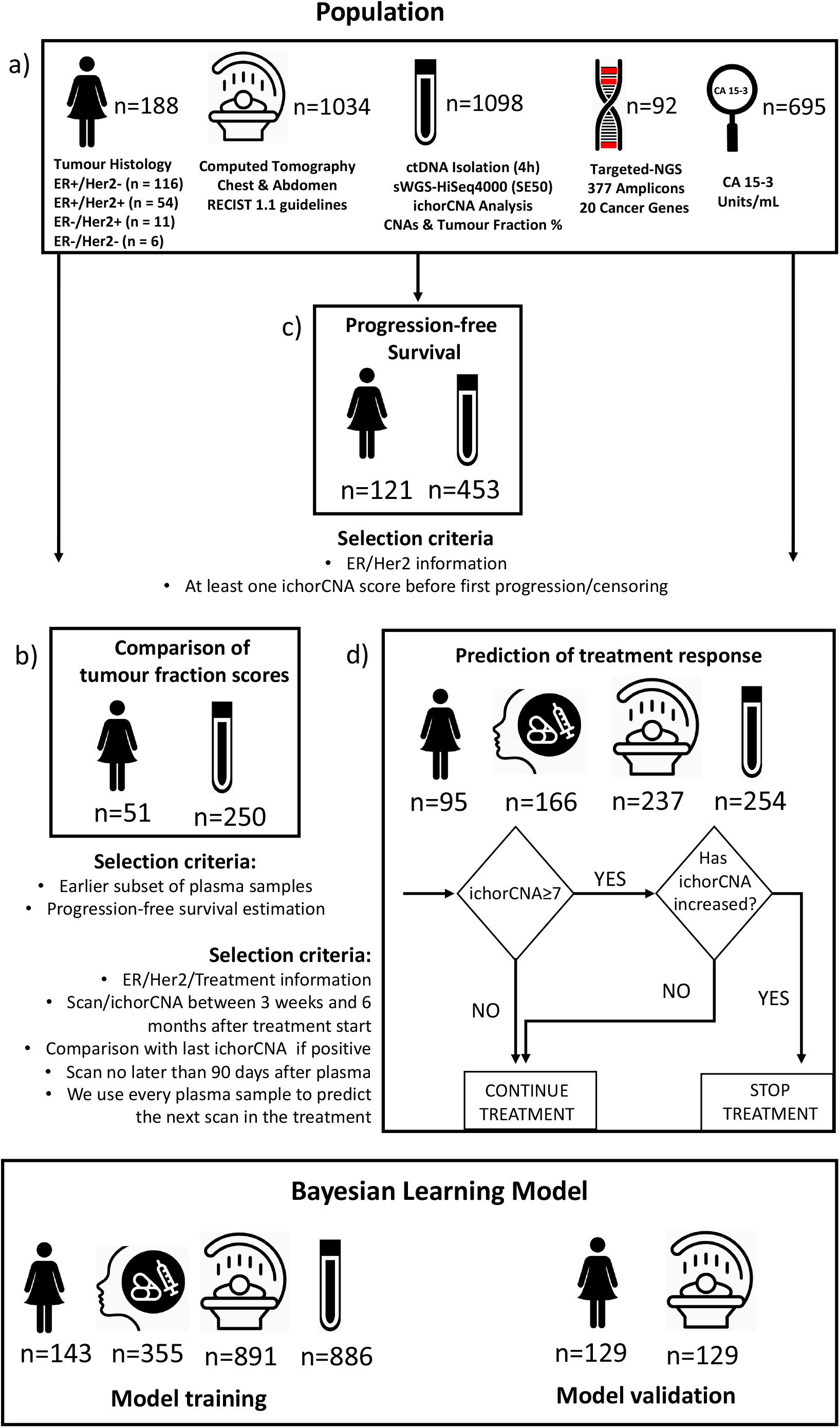
Clinical cohort and sample analysis with histology, sample timelines and treatment types. A) Patient numbers, histology and sample processing. B). Timeline of ctDNA collection per patient C) Treatment types assessed. *one patient was excluded as no relevant CT staging scan. 29 samples were not used for sWGS library preparation due to the very low levels of DNA in these samples (<5ng DNA total).

### Sample Processing and analysis

In total 1,098 blood samples were collected in ethylenediaminetetraacetic acid (EDTA) tubes and processed within 1 hour for plasma and buffy coat separation (Supplementary Methods). DNA was extracted from plasma and buffy coat and sequencing libraries for sWGS were prepared using 5 ng of cfDNA from each sample and 50 ng of DNA from buffy coat using the ThruPLEX® Tag-seq Kit (Takara Bio, Inc., Shiga, Japan) as described in the manufacturer’s instructions. The sequencing libraries were purified and quantified as detailed in the Supplementary Methods.

Targeted sequencing of 20 breast cancer specific genes (NGTAS), as described in Gao, *et. al*.^14^, was also performed using 5ng DNA and library preparation as above for sWGS. Samples were then amplified in triplicate with specific primers using the Fluidigm Access array™ platform.

All libraries were sequenced using an Illumina HiSeq 2000, at a mean depth of 0.1x for sWGS and >100x for NGTAS.

Serum Ca15-3 levels collected as part of routine clinical care were analyzed at the Cambridge University Hospitals biochemistry laboratory (accredited by the United Kingdom Accreditation Service).

### Bioinformatic analysis

Sequencing data was processed and analyzed as described in the Supplementary Methods. The copy number profiles produced using QDNAseq^15^ were used as input into three algorithms for tumor fraction estimation (ichorCNA^11^, z-score^12^ and t-MAD^13^, see Supplementary Methods). Mutational profiling was performed using the NGTAS pipeline^14^, with the maximum variant allele frequency (VAF) of any somatic mutation detected used for assessing its predictive performance.

### RECIST Criteria

Progression-free survival (PFS) for each line of treatment was calculated using RECIST 1.1^16^ guidelines to determine Progression Free Survival 2 (PFS2) as described previously (the time from the start of a line of treatment until objective progression on medical imaging using computed tomography (CT) of the chest and abdomen, or death)^17,18^. For each line of treatment, the CT scan prior to the start of this line of treatment was used as baseline. Imaging of the head was not included in the assessment of progression in this study. Longitudinal data available for two representative patients is shown in Figure 2.

**FIGURE 2:**
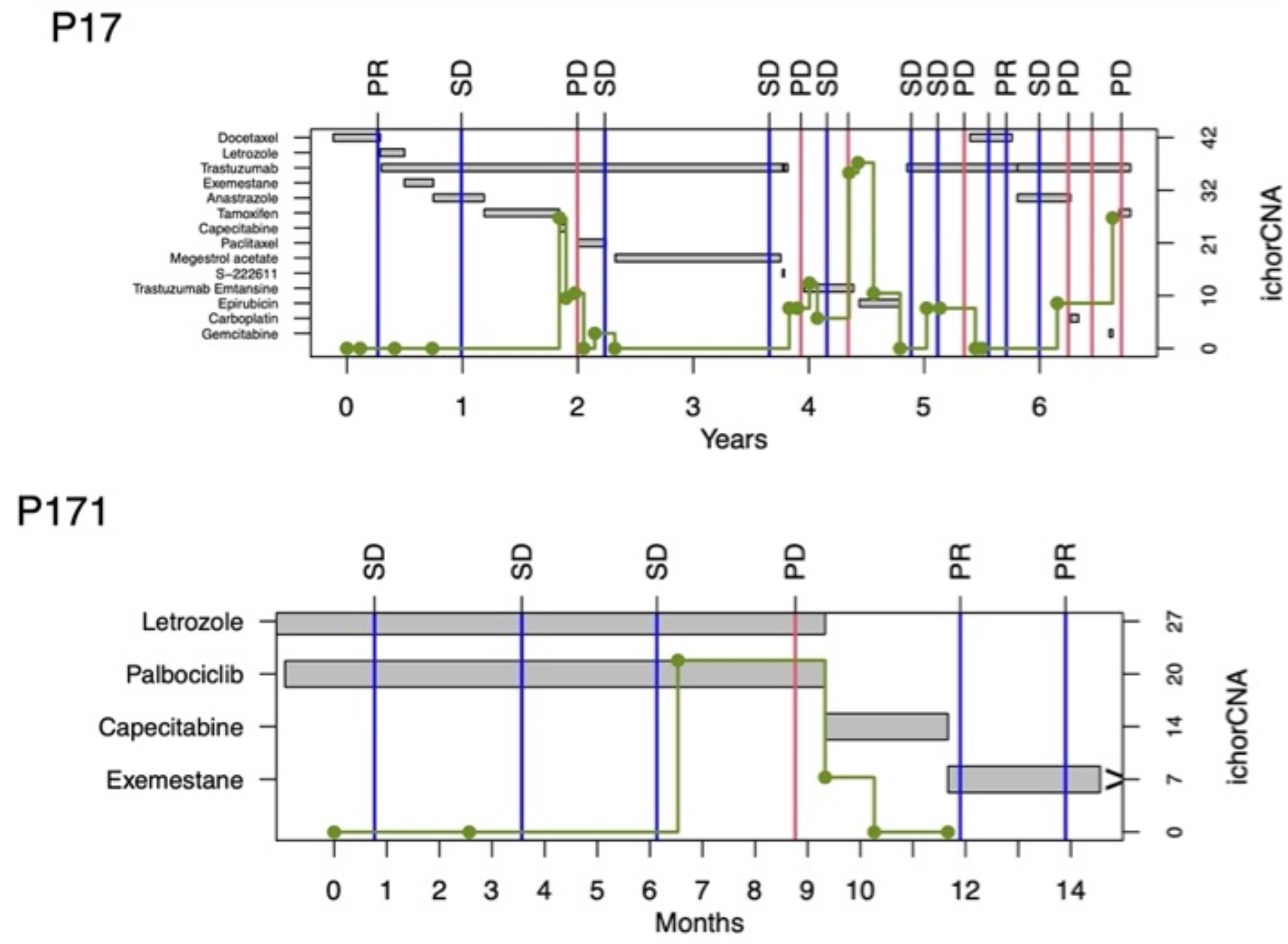
Profiles of two patients, showing the complexity of the longitudinal data available for each patient. Treatment regimes, CT Scans (PD=Progressive disease, SD=Stable disease, PR=Partial response) and ichorCNA scores are shown. Treatments ongoing are labelled with a > symbol.

## RESULTS

### ichorCNA best predicts progression-free survival

Several methods for ctDNA fraction estimation using sWGS CNA data of DNA extracted from plasma have been proposed, including ichorCNA^11^, z-scores^12^ or t-MAD^13^. In order to identify which method performed best at estimating tumor burden in plasma, we used a discovery dataset consisting of the first 51 patients totaling 250 plasma samples. We built univariable time-dependent Cox models for PFS using each measure (ichorCNA, t-MAD, or z-score) individually. Using the c-index^20^ to assess the three models showed ichorCNA performed best (ichorCNA c-index=0.71, se=0.05; t-MAD c-index=0.68, se=0.06; z.OR c-index=0.61, se=0.06).

We subsequently determined a threshold to identify patients at high risk of progression in order to facilitate clinical implementation. We used a spline term to model the effect of ichorCNA score on the hazard of progression and then fitted a segmented linear regression. Using a subsample of the patients (n=70, Supplementary Methods) this revealed a linear increase in the risk of progression followed by a changepoint at a mean score of 6.7% (Supplementary Figure 1 panel a, n=70). As expected, this categorization of the score showed lower predictive power than the continuous model in the subset of samples not selected in each iteration (Supplementary Methods, mean c-index=0.64, vs. 0.71, n=51), though the introduction of a static threshold makes clinical implementation easier. By choosing a threshold of 7%, the hazard ratio of progression for high ichorCNA score (≥7%) was 5.97 [3.72, 9.59] for the whole cohort (n=121). When we stratified tumours by subtype, we observed differences in predictive ability. In ER+HER2-patients, the hazard ratio was 5.60 [2.80, 11.18] and the expected time until progression for patients with a ‘low risk’ ichorCNA score (<7%) was 19.8 months, versus 8.2 months if the ichorCNA score was high. The prognostic effect was even higher for HER2+ patients, with a hazard ratio of 7.55 [3.39, 16.82] and a difference in the expected time of progression of 31.1 vs. 5.0 months. Figure 3 shows the predicted survival curves for two patients with high and low ichorCNA scores.

**FIGURE 3:**
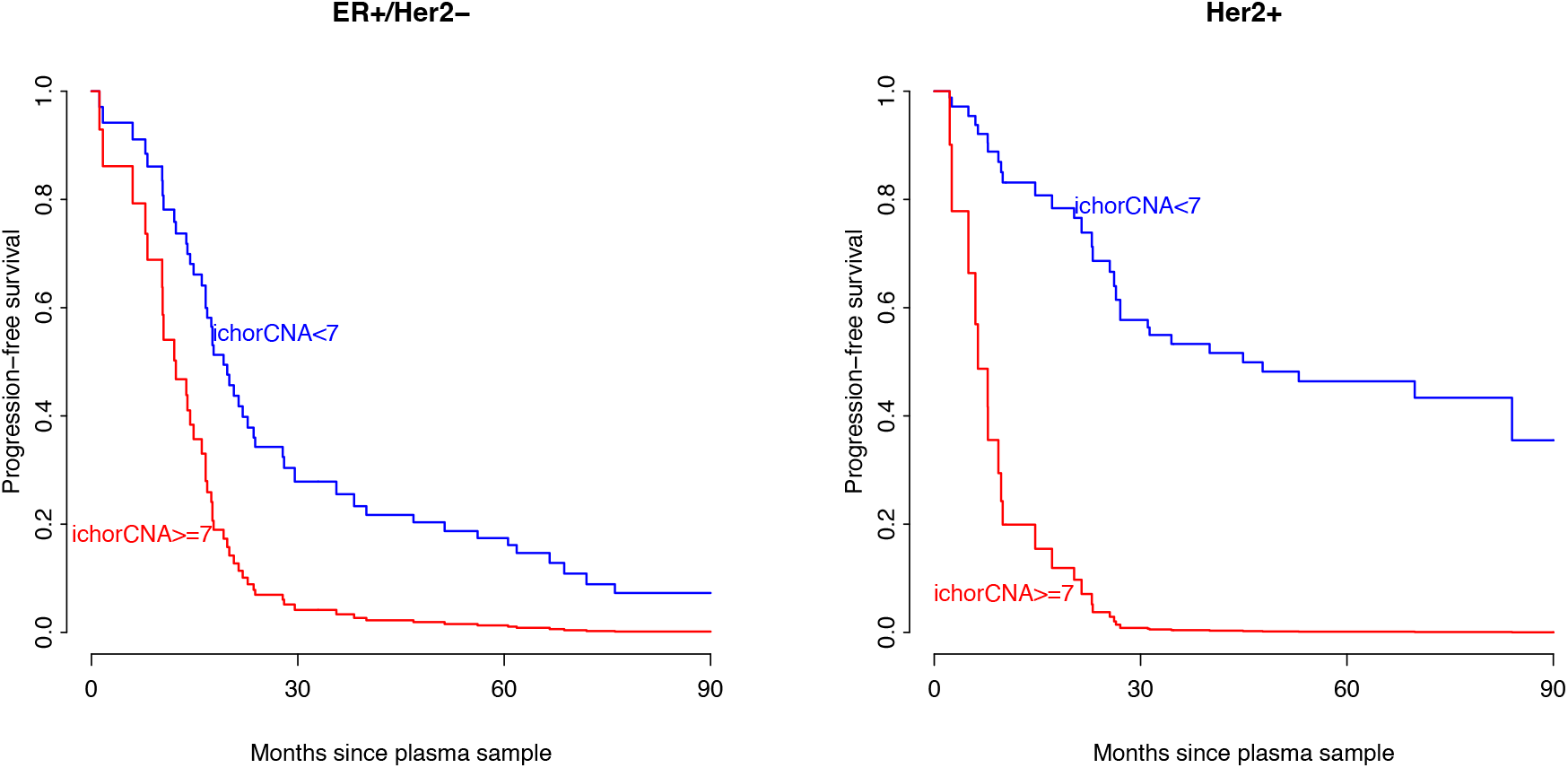
Predicted progression-free survival curves for two patients with low and high ichorCNA scores. The hazard ratio for each subtype has been obtained from a different model fitted for each disease subtype.

We observed that ichorCNA can also be used to predict overall survival. Using a continuous score with a linear term (p=9.77×10-8, hazard ratio: 1.08, [1.05, 1.11]) and the common threshold of 7%, a higher ichorCNA value was associated with an increased risk of death (p-value: 0.001, hazard ratio: 7.40, [2.38, 23.00]). This result highlighted the ability of ichorCNA to predict prognosis and the utility of our proposed threshold.

### Comparison with targeted sequencing (NGTAS) and CA15-3

Targeted mutational sequencing data was also available in 92 samples obtained from 22 patients (described in Gao *et. al*.^14^). In this smaller sub-cohort and using the maximum VAF observed in the sample, we did not observe a significant effect on the hazard of progression (p=0.24) while the ichorCNA remained significant (p= 0.034). Combining the two scores into the same model did not improve the fit (p=0.44).

We also looked at the ability of CA15-3 to predict response. For this analysis we compared data from a sub-cohort of 66 patients where 695 CA15-3 measurements were available with the corresponding ichorCNA values for the matched patient samples. Acknowledging the limitations of comparing both datasets (different number of patients and different sampling intervals, with the CA15-3 and the ichorCNA not always taken on the same day), a similar model using CA15-3 showed lower performance in terms of c-index (0.60, se=0.074) than the model using ichorCNA scores.

Considering a threshold of 0.01% VAF for NGTAS and 31U/ml (positivity threshold) for CA15-3, there was 78% concordance between NGTAS and ichorCNA, and a 60% concordance between ichorCNA and CA15-3. Figure 4 shows the instances where these measures showed discrepant results. Although the sampling times were different, comparing these values with the closest CT scan gave a better performance to the maximum VAF from NGTAS based on the area under the curve, but with a very small number of observations (0.78, n=15, ichorCNA=0.63, n=96 and CA15-3=0.59, n=81). Given these results, we decided to focus on ichorCNA for the rest of the study.

**FIGURE 4:**
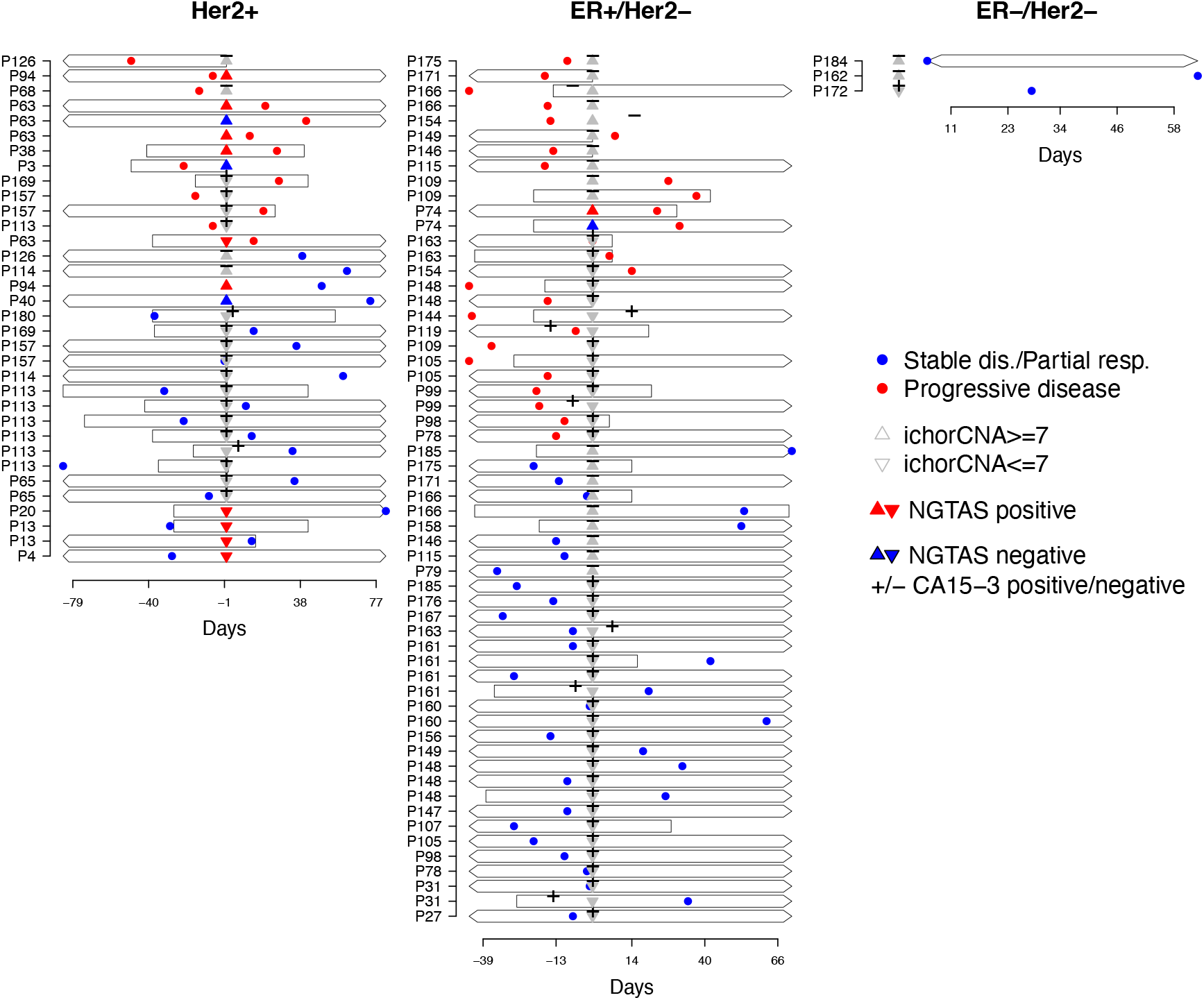
Discrepant results of ichorCNA measured with sWGS, mutant VAF measured with NGTAS and CA15-3. 96 instances where the CT Scan was done less than 90 days from or to the plasma sample and the CA15-3 was taken 15 days apart from the plasma sample measure are considered, Discrepancies are considered based on the 6% WHY 6% AND NOT 7%??threshold for ichorCNA, 30 for CA15-3 and 1% for VAF.

### ichorCNA ctDNA fraction predicts treatment response

Using the ichorCNA 7% threshold we evaluated its ability to predict subsequent response or resistance to treatment at each time point. We used the following rules: (i) for a prediction of response ichorCNA <7% or ≥7% if a decrease from the previous time point, (ii) for a prediction of progression ichorCNA ≥7% and an increase (or no change) from a previous time point (when available). The preceding time points needed to be on the same treatment to be relevant for decision-making. The application of these rules produced a sensitivity of 0.42 and a specificity of 0.90. The median time prior to prediction of progression for concordant decisions (stopping treatment) was 36 days (versus 29 days for discordant). For concordant decisions about continuing treatment the median time prior to the CT scan was 40 days (versus 38 days for discordant). These differences in the time where the decisions were made were not statistically significant and would not explain the difference in predictive ability.

### A Bayesian machine learning model (BAY-ML) to predict treatment response

Motivated by these findings, we developed a novel statistical model to predict treatment response (based on RECIST criteria) using the full history of ichorCNA scores and CT imaging (Figure 5a). The model comprises two components, one that includes the characteristics common to the cohort (ER/HER2 status and treatment regime) and another that models the patient specific longitudinal ctDNA scores and disease progression measurements on CT. The model is fitted using a two-stage approach: in the first stage, the evolution of the repeated ctDNA measurements is summarized by random effects obtained by fitting a linear mixed effects model, and in the second stage, the resulting random effects are used as covariates in a logistic regression to predict the risk of progression. Both steps include a set of independent variables, such as the treatment regime and the tumor (see Supplementary methods). The model adapts to each patient learning from common cohort’s effects such as the current treatment or the tumor subtype in both the ichorCNA trajectories and the probability of progression, but the model also learns from specific features of the patient. Leaving the final observation out for each patient (see Supplementary methods for details), we evaluated the sensitivity for predicting progressive disease when using ctDNA information at several clinically relevant specificity thresholds (Table 1). Leaving the last observation out in the model estimation, at 66% specificity, the sensitivity for detecting PD was 75%, significantly higher than our previous model based on a simple stopping rule. Figure 5b shows the receiver operator characteristic (ROC) curve for the model, highlighting the improvement in predictive performance, particularly in sensitivity, when iterative ichorCNA data is included.

**FIGURE 5:**
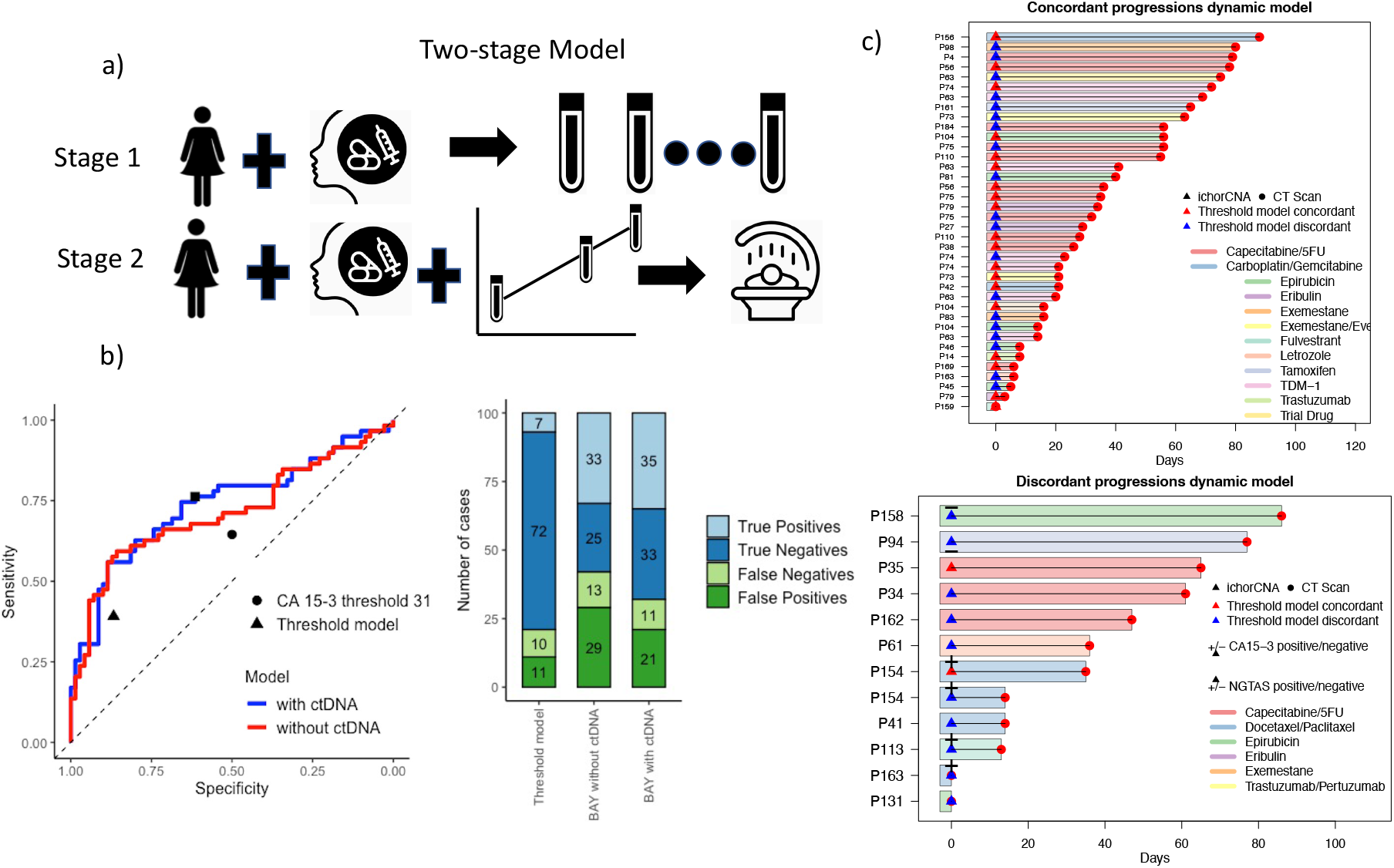
BAY-ML model. A) Visual summary of the two-stage model. B) Left: Receiver operator characteristic (ROC) curve of our dynamic predictive model. The predictions were obtained with the last CT scan for each patient, left out when the model was fit. Right: Number of true/false positives and negatives over 100 patients when the simplest threshold model and when the longitudinal ctDNA scores are considered or not into the BAY-ML model. C) Instances where the model correctly predicted progression and instances where it did not, comparing the available information at that moment.

**TABLE 1.** Sensitivity at various levels of specificity (last observation not included for each patient)

|  | Specificity |  |  |  |
| --- | --- | --- | --- | --- |
|  | 71% | 69% | **66%** | 61% |
| Sensitivity | 68% | 69% | **75%** | 76% |

### Treatment types

We looked at the predictive capability of ichorCNA for different treatment types. Predictions of response to targeted therapy (mainly CDK4/6 inhibitors and anti-HER2 therapy) and chemotherapy showed a higher concordance with CT results than endocrine treatment alone though this would need to be substantiated with larger data sets for statistical significance (see Figure 6). Blood test sampling was also less frequent for patients on endocrine treatment alone which could also explain this apparent difference.

**FIGURE 6:**
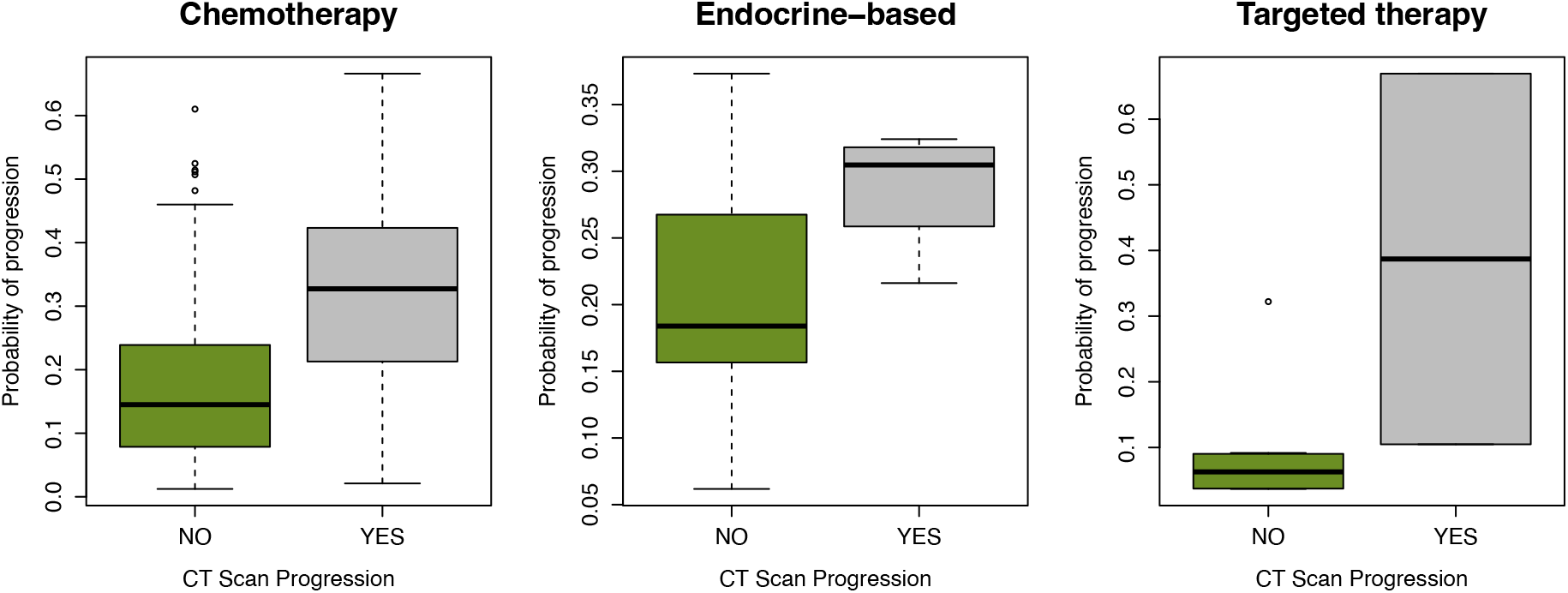
Analysis showing the association between the probability of predicting progressive disease and the result of the CT scan depending on different treatment types.

### Discordant result analysis

In order to understand better the limitations of ichorCNA we evaluated decisions whereby the ichorCNA values did not agree with disease status as measured using RECIST 1.1.

The ichorCNA score was high (predictive of treatment failure) but the subsequent CT showed RECIST stable disease in 24 instances. In 11/24 cases (46%), progressive disease was observed on the subsequent CT scan, suggesting early detection of progression by ctDNA, with a median lead time of 4.8 months. In 2/24 cases disease progression was observed in the brain (not evaluated as part of the RECIST criteria for this study).

In the cases where progression occurred on a CT scan but were not detected by ichorCNA (22 instances in total), in 5 of these cases this followed treatment with palliative radiotherapy which we hypothesize may have led to a reduction in ctDNA levels, separate to other systemic treatments. In 2/22 of cases a mixed response was seen, and it may be that having additional information from ctDNA could have aided the decision-making process. For an additional 6/22 cases low volume changes were seen (<10mm in <2 lesions) which may have had limited clinical impact. In 60% of these cases clinical management continued unchanged.

## DISCUSSION

We have demonstrated the real-world performance of ichorCNA in predicting treatment response for metastatic breast cancer patients and have used a novel Bayesian machine learning approach which uses longitudinal data, improving its predictive capability. This approach has several advantages over other methodologies. Firstly, all data is useful (including ichorCNA = zero) as it evaluates changes across the whole genome, having an important advantage over targeted sequencing, whereby a lack of somatic mutations in ctDNA only means that those specific mutations have not been detected. Moreover, within our study the average turnaround time, including library preparation and sequencing, was less than one week, and the assay we used was cheaper than currently available alternatives, with a typical cost of £100 per sample including processing costs. This compares favourably to commercially available ctDNA tests e.g: FoundationOne Liquid CDx (300 gene-panel analysis at ~£5,000 per sample; MSK-IMPACT™ (468 gene-panel analysis at ~£2,000-3,000 per sample); Both tests offer an outcome from the analysis in 2 weeks turnaround time what is twice as long than our in-house test. In addition, ctDNA monitoring could reduce the need and frequency of CT scanning if levels continued to be low.

Altogether, this shows that the method we describe in this study can be cheaply deployed within the clinic for therapy monitoring in real time. Importantly, the prediction capability was agnostic of breast cancer subtype and treatment regime, though it may be that predicting response to targeted therapy and chemotherapy may be slightly more reliable than for endocrine treatment alone. This may reflect the biology of disease as patients on targeted treatment and chemotherapy are likely to have more aggressive disease or have a higher tumor burden potentially making ctDNA levels higher and more dynamic. Moreover, cytostatic treatments are probably less likely to cause large changes in ctDNA compared with cytotoxic treatments. For a subset of cases, we also had data on CA15-3 levels and somatic mutations from a targeted sequencing panel. From this analysis ichorCNA was more accurate than CA15-3, and targeted sequencing data did not improve the predictive power with the caveat these comparisons were in smaller numbers of patients.

Due to lower patient numbers, we were less able to comment on any specifics for triple negative breast cancer. A prior study in metastatic triple negative breast cancer patients found a similar link between PFS and ichorCNA score^19^ and we do not believe there would be any significant difference for this subtype though the optimal threshold may be different. This also highlights the potential benefits of using machine learning to improve predictive power. We initially developed a simple stopping rule that is easy to implement in the clinic that is prognostic of progression and can predict treatment response with moderate success. We improved on this using a dynamic model that uses the history of the patient to predict more accurately the probability of progression under a given treatment.

Thinking more broadly about estimating ctDNA it could be that this is a better marker of overall disease activity than purely measuring disease on a CT scan. In a recent publication evaluating the use of ctDNA in the neoadjuvant setting it was shown that the presence of residual ctDNA post operatively was more predictive of relapse than pathological complete response^20^. We hypothesize that using the approach we describe here could provide benefits in terms of both quality of life, by reducing unnecessary toxicity, and increased access to more efficacious treatments in a timely fashion. This is also a realistic prospect as genomic assessment using ctDNA is rapid and relatively inexpensive making it accessible to public health systems. Our findings need to be fully evaluated in a prospective randomized clinical trial to assess the use of ctDNA to make treatment decisions, comparing it also to endpoints such as quality of life and overall survival.

## DATA AVAILABILITY

Supplementary Tables 1-5 contain the data used in this study.

## ACKNOWLEDGEMENTS

E.J.B was supported by the Cancer-ID European Consortium (SWAI-083), Academy of Medical Sciences (AMS-SGCL13-Beddows) & the Cambridge Breast Cancer Research Unit (CBCRU) as part of the Cancer Research UK Cambridge Centre [C9685/A25117]

M.O.D was supported by Servier Laboratories, France (RDAG/432) and by the Cancer Molecular Diagnostics Laboratory, Cambridge (Biomedical Campus).

O.M.R. was supported by the NIHR Cambridge Biomedical Research Centre (BRC-1215-20014) and the Medical Research Council (UK; MC_UU_00002/16).

S-J.S. was supported by a Wellcome Trust PhD Clinical Training Fellowship [grant number: 106566/Z/14/Z].

J.K. was supported by the Experimental Cancer Medicine Centre, Cambridge

J.L was supported by the Cancer Research UK Cambridge Centre

C.C. was supported by funding from CRUK [grant numbers: A16942, A17197, A27657, A29580], an NIHR Senior Investigator Award [grant number: NF-SI-0515-10090], and a European Research Council Advanced Award [Grant number 694620]. ERC grant also supported partly the salaries of M.O.D., S.-F.C. and M.C.

N.R. was supported by funding from Cancer Research UK (grant numbers A20240 and A29580)

R.W. was supported by the Austrian Science Fund (J-4025).

This research was supported by the NIHR Cambridge Biomedical Research Centre (BRC-1215-20014). The views expressed are those of the authors and not necessarily those of the NIHR or the Department of Health and Social Care.

We are grateful for the generosity of all the patients that donated samples for analysis; all the staff at the Cambridge Breast Cancer Research Unit for facilitating the collection and processing of samples.

NR is co-founder and officer of Inivata Ltd. Inivata had no role in the conceptualization or design of the clinical study, statistical analysis or decision to publish the manuscript.

C.C. is a member of AstraZeneca’s iMED External Science Panel and Illumina’s Scientific Advisory Board and a recipient of research grants (administered by the University of Cambridge) from Genentech, Roche, AstraZeneca and Servier.

